# Identification of glutamine synthetase as a novel contryphan-Bt binding protein by his-tag pull down

**DOI:** 10.1101/2020.12.10.419135

**Authors:** Penggang Han, Ying Cao, Xiandong Dai, Shangyi Liu, Chongxu Fan, Wenjian Wu, Jisheng Chen

## Abstract

**Background:** Contryphan-Bt is a D-tryptophan-containing disulfide-constrained decapeptide recently isolated from the venom of *Conus betulinus*. The molecular targets of contryphans are controversial, and the identification of its interacting proteins may be of great importance.

**Methods:** His-tag pull down assays were performed to investigate binding proteins of contryphan-Bt from rat brain lysate. Bt-Acp-[His]_6_, a contryphan-Bt derivative containing hexahistidine tag, was synthesized and used as the bait. As a control, Acp-[His]_6_ was used to exclude nonspecific bindings.

**Results:** Glutamine synthetase was identified as a potential contryphan-Bt binding protein by pull down assays and subsequent LC-MS/MS. The binding of contryphan-B to glutamine synthetase was confirmed and determined using microscale thermophoresis, with a Kd of 74.02 ± 2.8 μM. The binding did not affect glutamine synthetase activity, suggesting that the interaction site was distinct from the catalytic center.

**Conclusions:** Glutamine synthetase was identified as a novel contryphan-Bt binding protein. This is the first report that the conopeptide binds to an intracellular protein, therefore offering a new concept and methodology for developing peptide toxins.

**Key Contribution:** This is the first report that the conopeptide binds to an intracellular protein, therefore offering a new concept and methodology for developing peptide toxins.

## 1. Introduction

Conopeptides from *Conus* venom target a broad range of membrane proteins such as ion channels, transporters and GPCRs [1]; however, studies focus on their cellular interacting proteins are not available. Numerous bioactive peptides from marine sources interact with intracellular proteins and exhibit diverse pharmacological activities [2]. It’s interesting and wealthy to find out whether conopeptides could act though interacting with intracellular proteins.

Contryphans are kinds of unusual conopeptides, first isolated from the venom of piscivorous cone snail [3]. Up to now, 16 contryphans are characterized at the protein level (Table 1). Although comprising only 7-12 amino acid residues, contryphans are characterized by a high degree of post-translational modifications including N-terminal amide cyclization, a disulfide bond between two cysteines, proline hydroxylation, tryptophan bromination, leucine or tryptophan epimerization, C-terminal amidation, and Glu γ-carboxylation. Most contryphans share the conserved sequence motif CO(D-W or D-L)XPWC [4], providing an interesting scaffold for molecular design. Due to the constraints of ring cyclization by the S–S bridge, the proline residue close to the N-terminus undergoes *cis-trans* isomerization, and the *cis* isomer is more abundant. Other residues, especially the D-W/L also contribute to the *cis-trans* isomerization [5], which will lead to a later-eluting peak on the RP-HPLC for the minor *trans* conformer [6, 7].

**Table 1.**
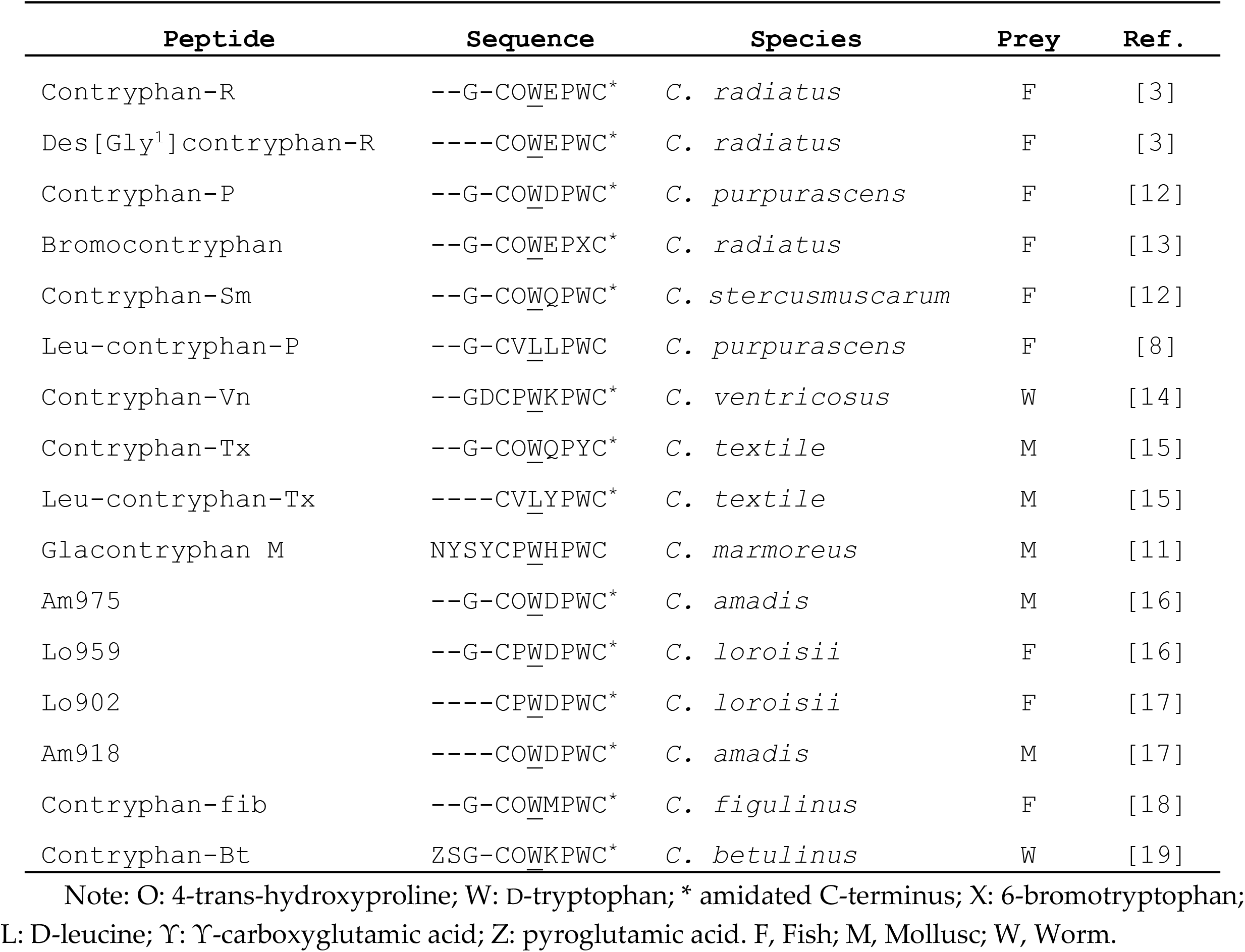
The contryphans identified at the protein level

Contryphans produce a “stiff-tail” syndrome in mice when intracranially injected, or triggers body tremor and mucous secretion syndromes when intramuscularly injected into fish [8, 9]. Some evidence indicates that contryphans might weakly act on voltage-gated calcium or potassium channels [10, 11]. However, the modulation is too weak (> 10 μM), and it’s unlikely that these channels are the physiological targets of contryphan [9]. The real molecular target of contryphans remains to be revealed.

Recently, we purified and characterized contryphan-Bt from *C. betulinus* venom [19]. Contryphan-Bt is a decapeptide that differs from other contryphans by an N-terminal pyroglutamic acid. The his-tag fusion protein system (his-tag pull down) is commonly used for the one-step purification of recombinant proteins and provides a convenient method to study protein-protein interactions [20]. Herein, we applied this strategy to search binding proteins of contryphan-Bt. Bt-Acp-[His]_6_, a derivative of contryphan-Bt with hexahistidine tag and aminocaproic acid (Acp) at the C-terminus, was synthesized and used as bait. Our results suggested that Glutamine synthetase (GS) from rat brain lysate is a contryphan-Bt-binding protein. The binding affinity was measured using microscale thermophoresis with a Kd of 74.02 ± 2.8 μM.

## 2. Results and discussion

### 2.1. Bt-Acp-[His]_6_ maintains the bioactivity of contryphan-Bt

To apply a his-tag pull down assay to identify proteins that associated with contryphan-Bt, Hexahistidine tag and spacer arm aminocaproic acid were introduced into the C-terminus of contryphan-Bt. The derivative was named Bt-Acp-[His]_6_ (Figure 1) and synthesized using Fmoc chemistry on Wang resin. Following chain assembly, the linear precursor was removed from the resin, and the disulfide bond Cys4–Cys10 was formed by air oxidation. The oxidized peptide was purified using RP-HPLC. Analytical HPLC detected a single peak, suggesting high purity (Figure 2). MALDI-TOF-MS analysis detected a 2122.95-Da ([M + H]^+^) ion (Figure 3), consistent with the calculated mass of Bt-Acp-[His]_6_ (2122.89 Da, [M + H]^+^). Acp-[His]_6_ was also synthesized and would be used as a control bait in pull down assay.

**Figure 1.**
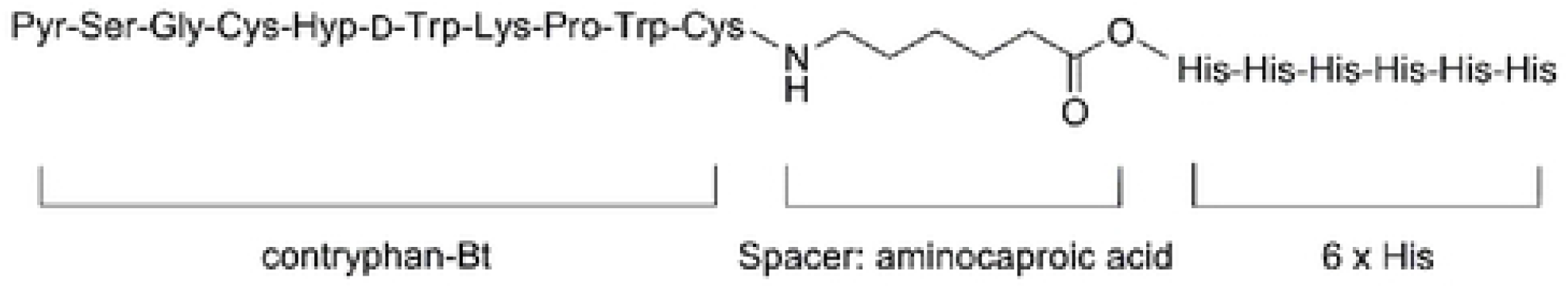
Structure of Bt-Acp-[His]_6_. Bt: contryphan-Bt; Acp: aminocaptoic acid; [His]_6_: his-tag composed of six histdine residues.

**Figure 2.**
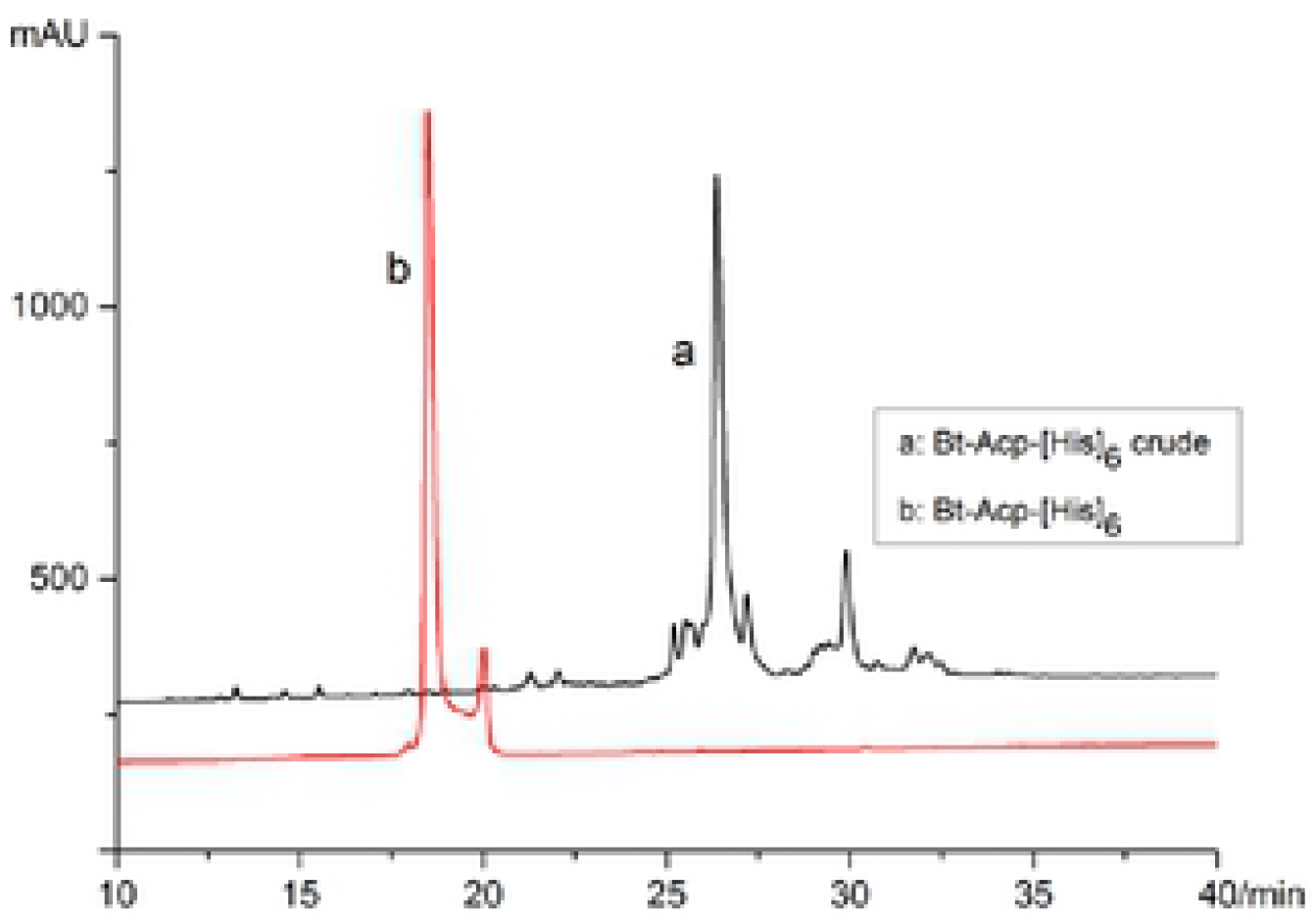
RP-HPCL profiles of crude and oxidized Bt-Acp[His]_6_ a: crude peptide; b: oxidized and purified peptide.

**Figure 3.**
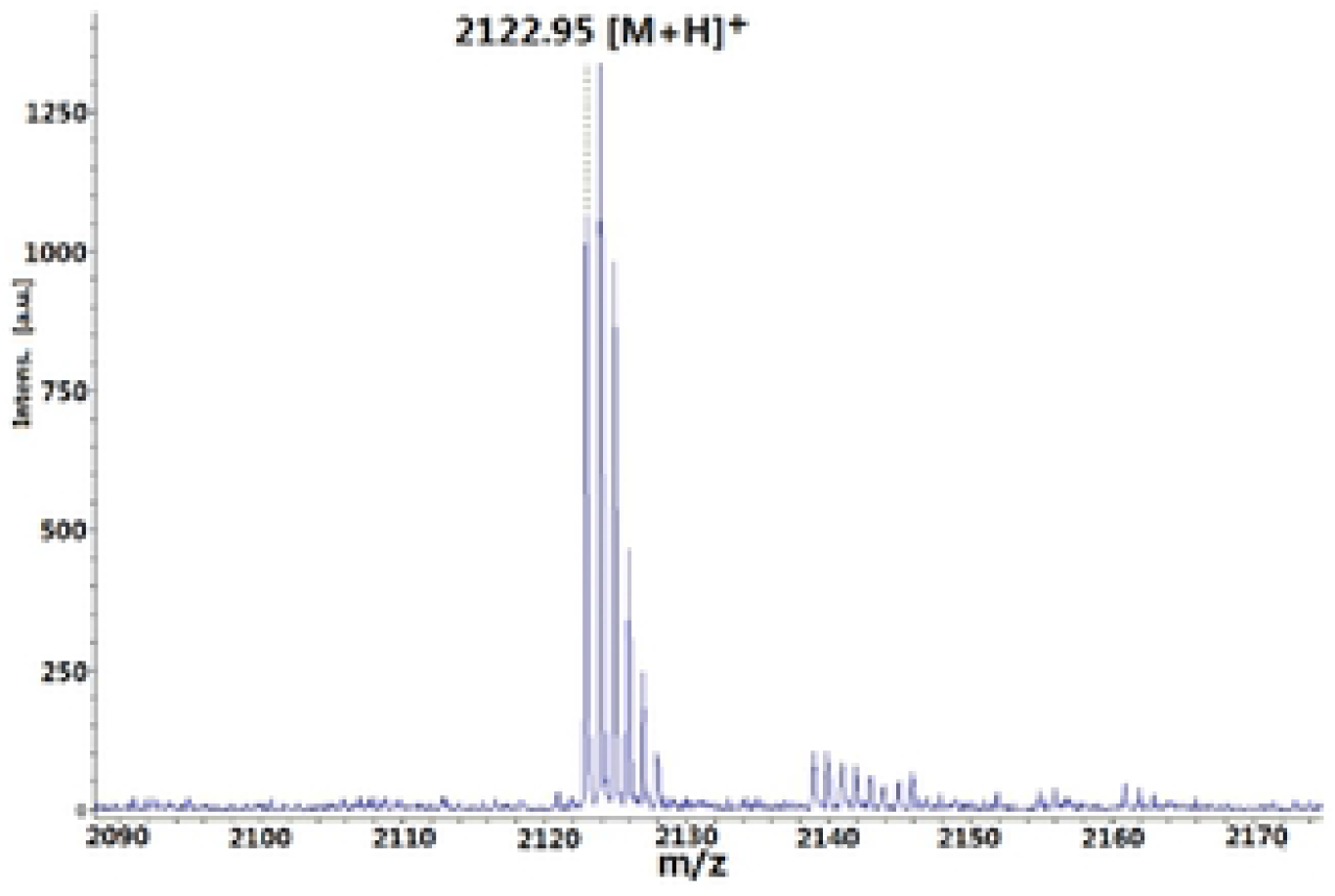
MALDI-TOF-MS analysis of Bt-Acp-[His]_6_([M + H]^+^.

To assess the effect of Acp and Hexahistidine tag on the bioactivity of contryphan-Bt, Bt-Acp-[His]_6_ was intracranially injected into mice as previously described [19]. Consistent with contryphan-Bt, characteristic symptoms including stiff-tail, body tremor, and barrel-rolling syndromes were observed, suggesting that the Acp and Hexahistidine tag modifications did not significantly alter the bioactivity.

### 2.2. Glutamine synthetase (GS) coelutes with Bt-Acp-[His]_6_ but not with Acp-[His]_6_

To search for cellular proteins that may interact with contryphan-Bt, pull down assays were performed with Bt-Acp-[His]_6_ and Acp-[His]_6_ as the baits, respectively. Acp-[His]_6_ was used as control to exclude nonspecific bindings. After incubation with rat brain lysate, samples were loaded onto charged cobalt spin columns. Nonspecifically bound proteins were removed using 10 mM imidazole solution. The bound proteins were then eluted with 350 mM imidazole and visualized by Coomassie Blue staining after SDS-PAGE. A ~42 kDa protein was specifically detected in Bt-Acp-[His]_6_ elution buffer (Figure 4a).

**Figure 4.**
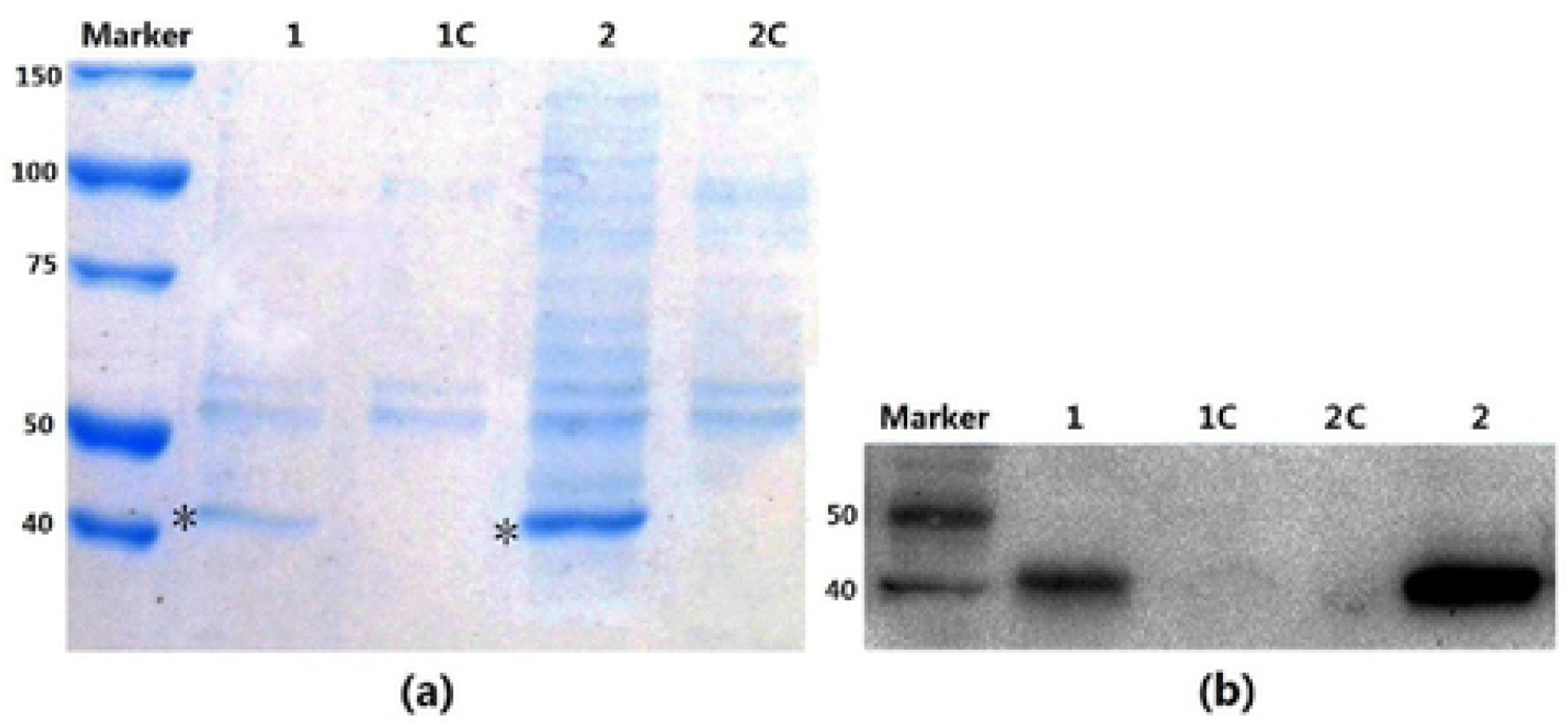
Glutamine synthetase (GS) coelutes with Bt-Acp-[His]_6_ but not with Acp-[His]_6_. Pull-Down assays using Bt-Acp-[His]_6_ or Acp-[His]_6_ (control) were conducted and the elution buffer were separated by10% SDS-PAGE. 1 and 2: elution buffer of Bt-Acp-[His]_6_; 1C and 2C: elution buffer of Acp-[His]_6_. Asterisks indicate the ~42 kDa protein detected specifically in the experimental group (a). The proteins were transferred to a PVDF membrane, and GS was detected using an anti-rat GS antibody (b).

The ~42 kDa protein band was manually excised and identified by LC-MS/MS. A search of the Mascot database suggested that the protein was glutamine synthetase (GS), with 48 matching peptides and 37.8% sequence coverage (Figure 5). In agreement with LC-MS/MS analysis, western blotting using anti-GS antibody only detected signals in the Bt-Acp-[His]_6_ lane (Figure 4b). The results indicate that GS coelutes with Bt-Acp-[His]_6_ but not with Acp-[His]_6_, suggesting that GS may interact with contryphan-Bt.

**Figure 5.**
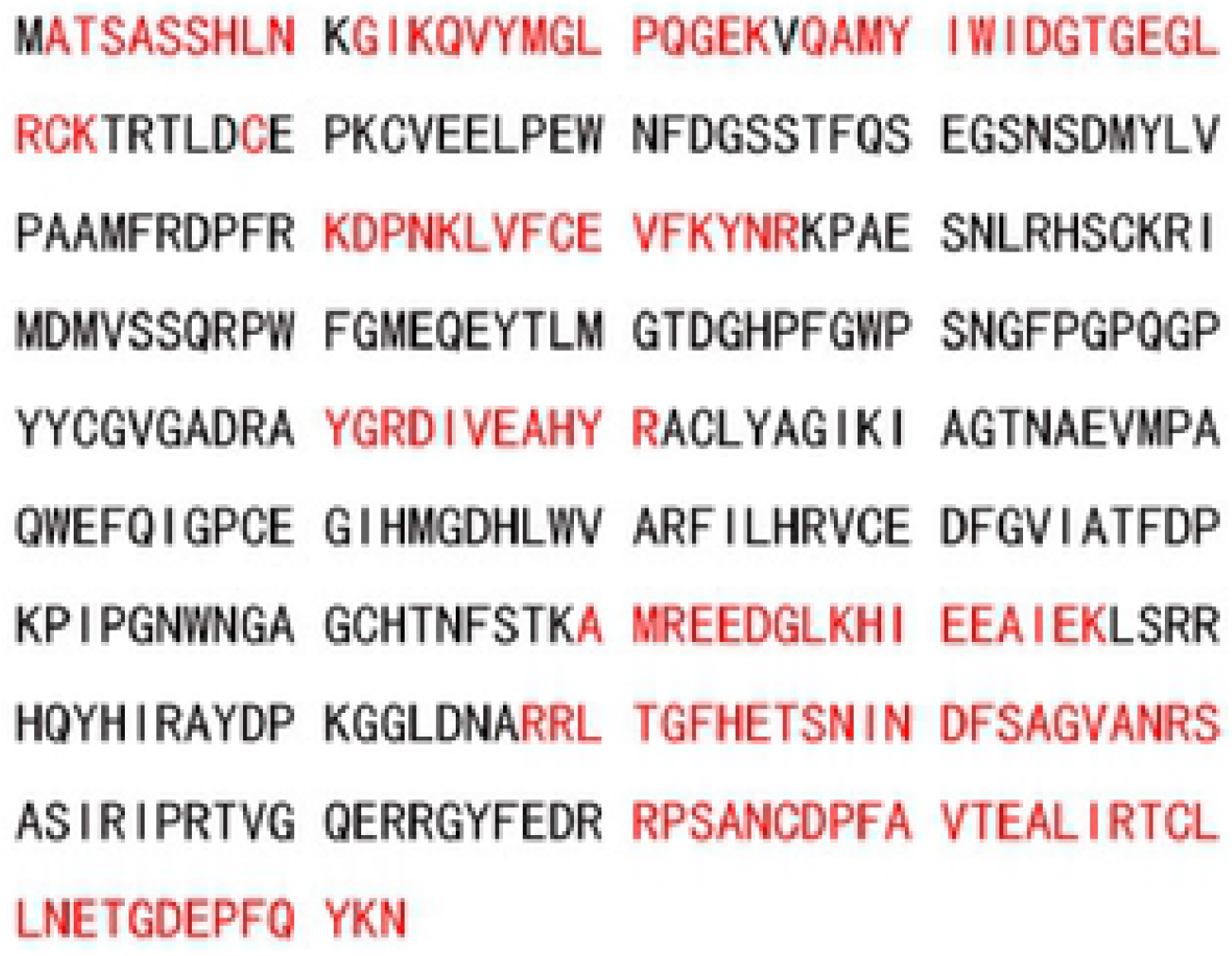
Amino acid sequence of rat Glutamine synthetase (373 residues). The peptides identified by LC-MS/MS are highlighted in red.

### 2.3. Contryphan-Bt directly binds to glytamine synthetase

To further validate the interaction between contryphan-Bt and glutamine synthetase (GS), we applied microscale thermophoresis (MST) to determine their binding affinity. MST is a powerful new method for the quantitative analysis of protein-protein interactions (PPIs) with low sample consumption. The technique measures the direct movement of molecules along a temperature gradient, an effect termed “thermophoresis” [21]. Thermophoresis depends on changes in molecular size, charge, or solvation shell. Binding of a nonfluorescent molecule to a fluorescently-labeled molecule changes at least one of these properties, which affects thermophoretic mobility [22]. Thus, MST could rapidly detect numerous and diverse biomolecular interactions under non-denaturing conditions.

Human recombinant GS was used for MST experiments, as it is commercially available and is well conserved to rat GS with an identity of 95.7%. The fluorescent dye NT-647-NHS -labeled GS was titrated with 15 serial dilutions of contryphan-Bt with concentrations varying from 0.5 mM to 32 nM. The resulting fluorescence signal was smooth, indicating the absence of aggregates. A concentration-dependent change in the thermophoresis of dye labeled GS was observed, suggesting a specific interaction between GS and contryphan-Bt. A sigmoidal binding curve was obtained by performing a Kd fit according to the law of mass action, yielding a Kd value of 74.02 ± 2.8 μM (Figure 6).

**Figure 6.**
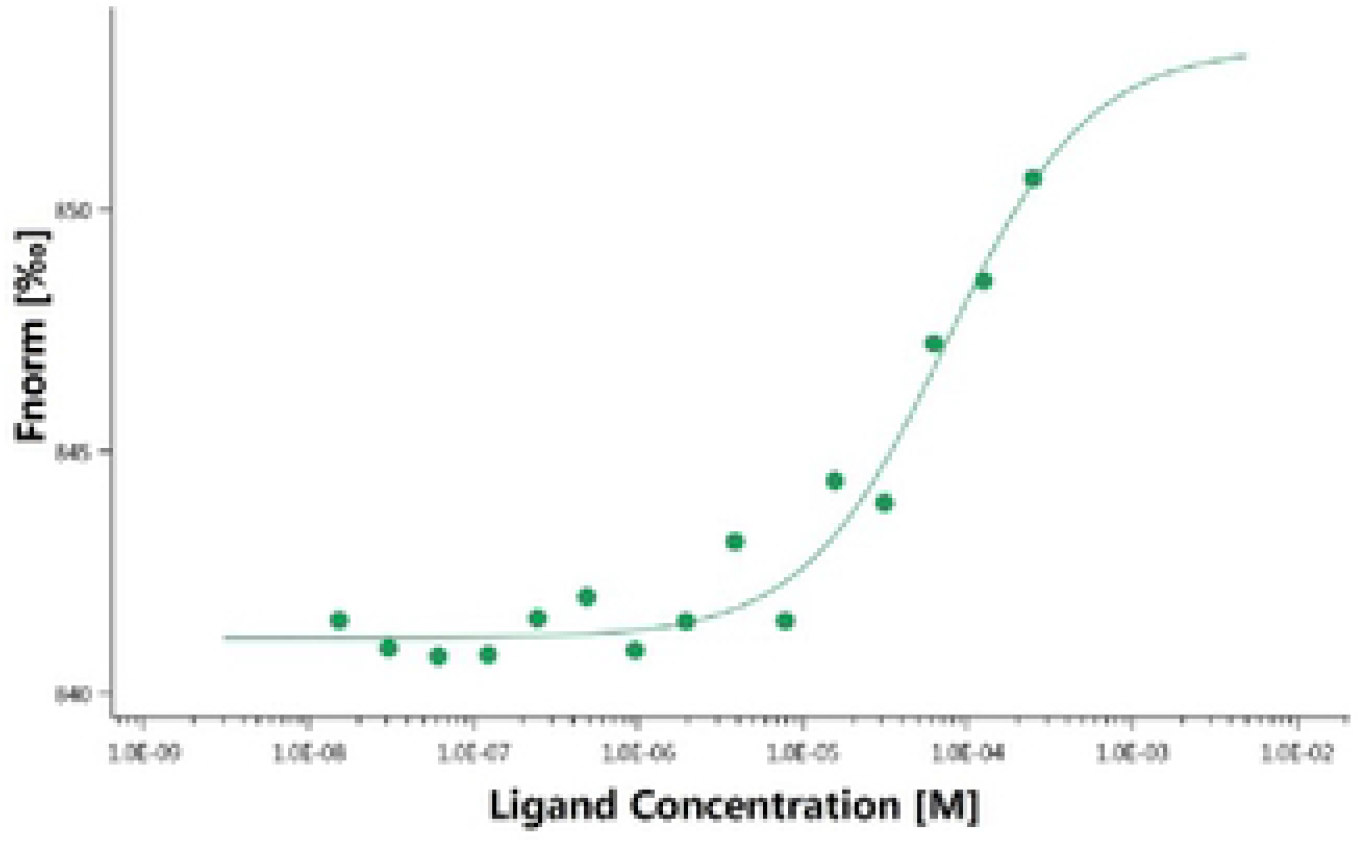
Dose-response curve generated by MST for the binding interaction between contrypgan-Bt and glutamine synthetase. The concentration of dye labeled GS was kept constant, while the contryphan-Bt concentration varies from 0.5nM to 32 nM. A Kd of 74.02 ± μM was calculated. Fnorm = normalized fuorescence.

### 2.4. Contryphan-Bt does not affect GS activity

Glutamine synthetase (GS) is a key enzyme, which catalyzes the ATP-dependent conversion of ammonia and glutamate to glutamine. In the brain, GS plays a central role in glutamate and glutamine homoeostasis [23, 24], in the termination of neurotransmission through glutamate, and in the detoxification of brain ammonia [25]. The GS activity can be assayed by the γ-glutamyl transferase reaction, in which γ-glutamyl-hydroxamate (γ-GH) is formed and gives a characteristic color reaction with ferric chloride (Figure 7) [26, 27].

**Figure 7.**
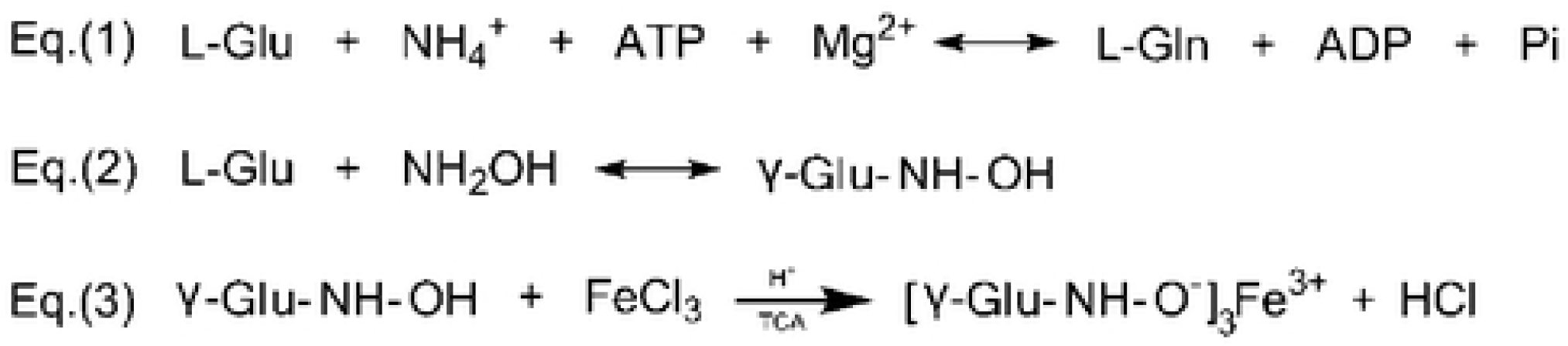
The γ-glutamyl transferase reactions for GS activity assay. Eq.(1): GS catalyzes the formation of L-Gln from L-Glu, driven by the hydrolysis of ATP. Eq.(2): When ammonia is replaced by hydroxylamine, γ-GH(γ-Glu-NH-OH) is produced. Eq.(3): γ-GH reacts with ferric chloride, yelding a colored which absorbs at 540nm.

To further evaluate the effect of contryphan-Bt on GS activity, GS from rat brain was partially purified using two-step acid precipitation method [28]. GS solution was incubated with contryphan-Bt with a range of 0-0.8 μM and subjected to activity assay. However, the absorbance at 540 nm showed no significant difference among the groups (data not shown), suggesting that binding of contryphan-Bt did not affect GS activity.

L-methionine sulfoximine (MSO) is a potent inhibitor of GS. In the presence of ATP and metal ions, MSO is phosphorylated by GS, and the phosphorylated form irreversibly binds to GS, permanently inactivating the enzyme [29]. Similarly, we assessed the effect of contryphan-Bt on the inhibition of GS by MSO. The results (data not shown) indicated that contryphan-Bt did not affect the inhibition of GS activity by MSO, suggesting that their binding sites on GS were distinct.

In summary, the data presented here proved that contryphan-Bt binds to GS but has no effect on the enzyme activity. Conopeptides were commonly considered to target membrane receptors. This is the first report that conopeptide binds to an intracellular protein, although the effect of the interaction remains to be defined. In another study, ConoCAP-a, a decapeptide isolated from *Conus Villepinii*, was found to produce a reduction of heart rate and blood pressure in rats by decreasing systolic calcium [30]. However, voltage-clamp experiments showed that conoCAP-a did not affect cell membrane channels and membrane receptors involved in the cardiovascular physiology, suggesting that conoCAP-a may require binding to an intracellular target protein that is essential for intracellular calcium handling. Membrane permeability assay indicated that conoCAP-a shows passive cell-permeability [30]. Together with our results, it was reasonable to suppose that some conopeptides may act though interacting with intracellular proteins.

## 3. Conclusion

In the study, a his-tag pull down strategy was adopted to investigate binding proteins of contryphan-Bt from brain lysates with Bt-Acp-[His]_6_ as the bait. Glutamine synthetase was identified as a contryphan-Bt-binding protein with a Kd of 74.02 ± 2.8 μM. To the best of our knowledge, this is the first report that conopeptide binds to an intracellular protein. Although the binding did not affect GS activity, our findings suggest that conopeptides, which were always thought to target membrane proteins, may interact with intracellular proteins as well.

## 4. Materials and methods

### 4.1 Materials

Rink amide resins, Fmoc-protected amino acids, and other reagents for peptide synthesis were acquired from GL Biochem (Shanghai, China). Gradient-grade acetonitrile (ACN), an anti-GS rabbit antibody, L-methionine sulfoximine, and protease inhibitor cocktail were obtained from Sigma-Aldrich (St. Louis, MO, USA). The BCA Protein Assay Kit, HisPur cobalt chromatography cartridges, MS-grade trypsin, goat anti-rat IgG-HRP, and western blot luminol reagent were purchased from Thermo Scientific (Waltham, MA, USA). Recombinant human GS was obtained from Prospec (Rehovot, Israel). All animal studies were in accordance with the NIH guide for the care and use of laboratory animals and were approved by the Beijing Institute of Technology Animal Institute Committee (SYXK-BIT-20181009001).

Automated peptide synthesis was conducted on a CEM Liberty peptide synthesizer (CEM, USA). RP-HPLC was performed using an Agilent 1100 system equipped with a dual-wavelength UV detector (Agilent, USA). MALDI-TOF-MS analysis was measured using a Bruker Ultraflex TOF/TOF mass spectrometer (Bruker Daltonics, USA) with α-cyano-4-hydroxycinnamic acid (CCA) as the matrix. LC-MS/MS data were acquired using a Q Exactive HF MS (Thermo Fisher Scientific) connected to an Easy-nLC 1200 nanoflow LC system (Thermo Fisher Scientific,). Microscale thermophoresis measurements were performed on a Monolith NT.115 (Nano Temper, Germany).

### 4.2 Peptide synthesis

Linear contryphan-Bt, Bt-Acp-[His]_6_, and Acp-[His]_6_ peptides were synthesized using standard Fmoc chemistry with Rink amide and Wang resins, respectively. The following side-chain protection groups were used: Fmoc-His(Trt)-OH, Fmoc-Gln(Trt)-OH, Fmoc-Ser(OBut)-OH, Fmoc-Gly-OH, Fmoc-Cys(Trt)-OH, Fmoc-Pro-OH, Fmoc-Hyp(OBut)-OH, Fmoc-D-Trp(Boc)-OH, Fmoc-Trp(Boc)-OH, Fmoc-Lys(Boc)-OH and Fmoc-e-Acp-OH. Peptides were cleaved from the resin using TIS/TFA/H2O mixture (2.5:2.5:95), precipitated in cold diethyl ether, and washed 5-times with cold diethyl ether. Crude peptides were purified using preparative RP-HPLC with a linear gradient of 0%–50% buffer B (acetonitrile containing 0.1% TFA) in 50 min. Purified, reduced peptides were oxidized in 25 mM NH_4_HCO_3_, pH 8, and then purified using preparative RP-HPLC. The purities of the final products were examined by analytic RP-HPLC, and molecular masses were validated by MALDI-TOF-MS.

### 4.3 Peptide Biossay

The biological activities of contryphan-Bt and Bt-Acp-[His]_6_ were assessed by intracranial injection into 10–20-day-old mice as previously described [19]. Peptides were dissolved in normal saline solution and subsequently injected using a syringe fitted with a 29-gauge needle. Negative-control mice were injected with an equal volume of normal saline. After injection, the behaviors of the mice were observed.

### 4.4. Preparation of Rat brain homogenate

Female SD rats (4-weeks-old) were provided a standard laboratory chow diet and water before the experiments. Rats were anesthetized using diethylether and then decapitated. The brains were immediately harvested, cut into small pieces, and homogenized on ice in 10 volumes of lysis buffer (50 mM Tris-HCl, pH 7.4, 150 mM NaCl, 1× protease inhibitor cocktail) using a glass homogenizer. The homogenate was centrifuged twice for 30 min at 4 °C and 6 000 g. The resultant supernatant was used fresh or stored at −80 °C. The total protein concentration (6.5 mg/ml) was determined using the BCA assay.

### 4.5. Crude extract of Glutamine synthetase from rats brain

Glutamine synthetase (GS) was partially purified from the rat brains using a modification of the method originally developed by Yamamoto [28]. Cooled acetone (10 volumes) was slowly added to the homogenate with constant stirring for 30 min. After filtration, the precipitate was washed twice with 10 volumes of cooled acetone, and a brain acetone powder was obtained by desiccation under high vacuum overnight. The acetone powder (1.5 g) was stirred for 30 min with 20 volumes of 5 mM mercaptoethanol and 150 mM KCl (pH 7.2), and then centrifuged at 11000 g at 4 °C for 40 min. The supernatant was placed in an ice bath, and the pH was adjusted to 6.2 by drop-wise addition of 1 M acetic acid with continuous stirring. After centrifugation, the precipitate was discarded, and the pH of the supernatant was slowly lowered to 4.2 using 1 M acetic acid. The solution was then centrifuged at 15000 g at 4 °C for 30 min, and the precipitate was extracted by stirring for 1 h with 0.1 M potassium phosphate buffer, pH 7.5, 5 mM mercaptoethanol, and 0.1 M glutamine. Following centrifugation at 15000 g at 4 °C for 30 min, the supernatant was used as the crude GS preparation.

### 4.6. GS assay and binding experiments

GS activity was measured using the forward reaction assay as previously described [26] with slight modifications. 50 μL contryphan-Bt solution, with increased concentration from 0– 2 μM (0, 0.2 μM, 0.4 μM, 0.8 μM contryphan-Bt), were added to 50 μL GS enzyme solution and incubated 30 min at 37 °C. The solution (100 μL) was then mixed with assay mixture (400 μL) comprising 100 mM imidazole chloride (pH 6.8), 50 mM L-Glu, 15 mM hydroxylamine (NH_2_OH), 30 mM MgCl_2_, and 7.5 mM ATP.guide After incubation at 37 °C for 20 min, the enzyme reaction was terminated by adding 1000 μL of a solution of 37 mM FeCl_3_, 10 mM TCA, 67 mM HCl. Insoluble material was removed by centrifugation, and the absorbance of the supernatants was measured at 540 nm in a spectrophotometer. The absorbance values represent the GS activity. Each assay was repeated three times.

### 4.7. Pull down assay

Bt-Acp-[His]_6_ was suspended in the brain lysate at 25 μM and incubated at 4 °C for 2 h with gentle stirring. The mixture was filtered (0.45 μm) and loaded onto an equilibrated cobalt spin column at 0.2 ml/ml. Nonspecifically bound proteins were removed using 15 volumes of wash buffer (50 mM PB, pH 7.4, 150 mM NaCl, 10 mM imidazole). Bt-Acp-[His]_6_ and its prey protein were eluted with 50 mM PB, pH 7.4, 150 mM NaCl, and 350 mM imidazole. The wash and elution effluents were collected. For evaluating nonspecific binding of proteins to the affinity matrix, we prepared Acp-[His]_6_ as a negative control, which was similarly incubated with the brain lysate, applied to the cobalt spin column and eluted.

### 4.8. LC-MS/MS analysis

Eluted samples were subjected to 10% SDS-PAGE followed by Coomassie Blue staining. The target protein band was excised from the gel, reduced in 10 mM DTT, alkylated with 50 mM IAA, and digested in the gel slices overnight using 25 μg/mL trypsin. The extracted peptides were loaded onto a silica capillary column (100 μm × 3 cm) packed with C18 reverse-phase resin (particle size, 3 μm; pore size, 150 Å). After desalting, peptides were separated using an analytical C18 column (75 μm × 15 cm, 3 μm, 150 Å) with a linear gradient of 5%–35% Mobile Phase B (acetonitrile containing 0.1% formic acid) at 600 nL/min for 70 min. The LC eluent was coupled to a Q Exactive HF MS. Mass spectra were acquired in positive-ion mode with automated data-dependent MS/MS on the five most intense ions detected in preliminary MS scans. MS/MS raw files were used to query the Swiss-Prot database using Mascot 2.3.

### 4.9. Western blotting

Eluted proteins were separated using 10% SDS-PAGE and electrophoretically transferred to a PVDF membrane. The membrane was blocked with 10% nonfat milk in TBST for 1 h at room temperature and then incubated with a rabbit anti-GS antibody overnight at 4 °C. HRP-conjugated anti-rabbit IgG was used as the second antibody. Protein bands were visualized using ECL reagents on an Amersham Imager 600 chemiluminescence system.

### 4.10. Microscale thermophoresis

Microscale thermophoresis (MST) experiments were conducted on a NanoTemper Monolith NT.115 system (NanoTemper, Germany). Recombinant human glutamine synthetase (GS) was labeled using a Monolith NT™ Protein Labeling Kit. GS was diluted to about 200 nM in label buffer. NT-647-NHS reactive dye was diluted in label buffer to a final concentration of 400 nM. 100 μL of protein was mixed with 100 μL of dye. The reaction mixture was incubated for 30 mins at room temperature in the dark. After labeling, unreacted free dye was eliminated and the solution was substituted to PBS (10 mM Na_2_HPO_4_, 2 mM NaH_2_PO_4_, 4.7 mM KCl, 135 mM NaCl, pH 7.4). A two-fold dilution series of contryphan-Bt was prepared in PBS, with the final concentrations ranging from 0.5 mM to 32 nM. 10 μL of labeled GS and 10 μL of dilutions of contryphan-Bt were mixed by pipetting up and down. Following incubation in the dark for 30 mins, the samples were transferred into Monolith NT standard treated glass capillaries and were subsequently subjected to MST analysis. All measurements were replicated three times at 20% LED power and 40% MST power at 25 °C. Data analysis, curve fitting, and quantification of the dissociation constant (Kd) were performed using NT Analysis software 2.2.4.

## Author Contributions

Conceptualization, Chongxu Fan and Jisheng Chen; methodology, Penggang Han, Shangyi Liu and Ying Cao; investigation, Penggang Han and Xiandong Dai; writing—original draft preparation, Penggang Han; writing—review and editing, Penggang Han, Chongxu Fan and Wenjian Wu; visualization, Penggang Han; supervision, Chongxu Fan; All authors have read and agreed to the published version of the manuscript.

## Funding

This research received no external funding.

## Conflicts of Interest

The authors declare no conflict of interest.

